# Integrating planar polarity and tissue mechanics in computational models of epithelial morphogenesis

**DOI:** 10.1101/138172

**Authors:** Katherine H. Fisher, David Strutt, Alexander G. Fletcher

## Abstract

Cells in many epithelial tissues are polarised orthogonally to their apicobasal axis. Such planar polarity ensures that tissue shape and structure are properly organised. Disruption of planar polarity can result in developmental defects such as failed neural tube closure and cleft palette. Recent advances in molecular and live-imaging techniques have implicated both secreted morphogens and mechanical forces as orienting cues for planar polarisation. Components of planar polarity pathways act upstream of cytoskeletal effectors, which can alter cell mechanics in a polarised manner. The study of cell polarisation thus provides a system for dissecting the interplay between chemical and mechanical signals in development. Here, we discuss how different computational models have contributed to our understanding of the mechanisms underlying planar polarity in animal tissues, focusing on recent efforts to integrate cell signalling and tissue mechanics. We conclude by discussing ways in which computational models could be improved to further our understanding of how planar polarity and tissue mechanics are coordinated during development.

## Introduction

A central problem in developmental biology is to understand how tissues form and repair in a highly reproducible manner. Key signalling molecules are spatially coordinated to provide positional information in developing tissues. While it has long been known that cells can sense and interpret such chemical gradients during pattern formation [1], mechanical forces are now recognised to also play a vital role in shaping tissues [2,3]. Increasing evidence suggests that these chemical and physical mechanisms are interconnected [4].

Morphogenesis is frequently driven by the dynamics of epithelial tissues, which line the majority of organs in the body. As well as being characterised by polarity along an apicobasal axis, epithelia often exhibit planar polarity orthogonally through the plane of the tissue (Fig. 1A) [5]. While it is possible for individual cells to become planar polarised, animal epithelial cells locally coordinate their polarity via intercellular transmembrane complexes (Fig. 1B) [6,7] to robustly generate uniform polarity across tissues, even when a global polarising signal is weak or noisy [8,9].

**Figure 1.**
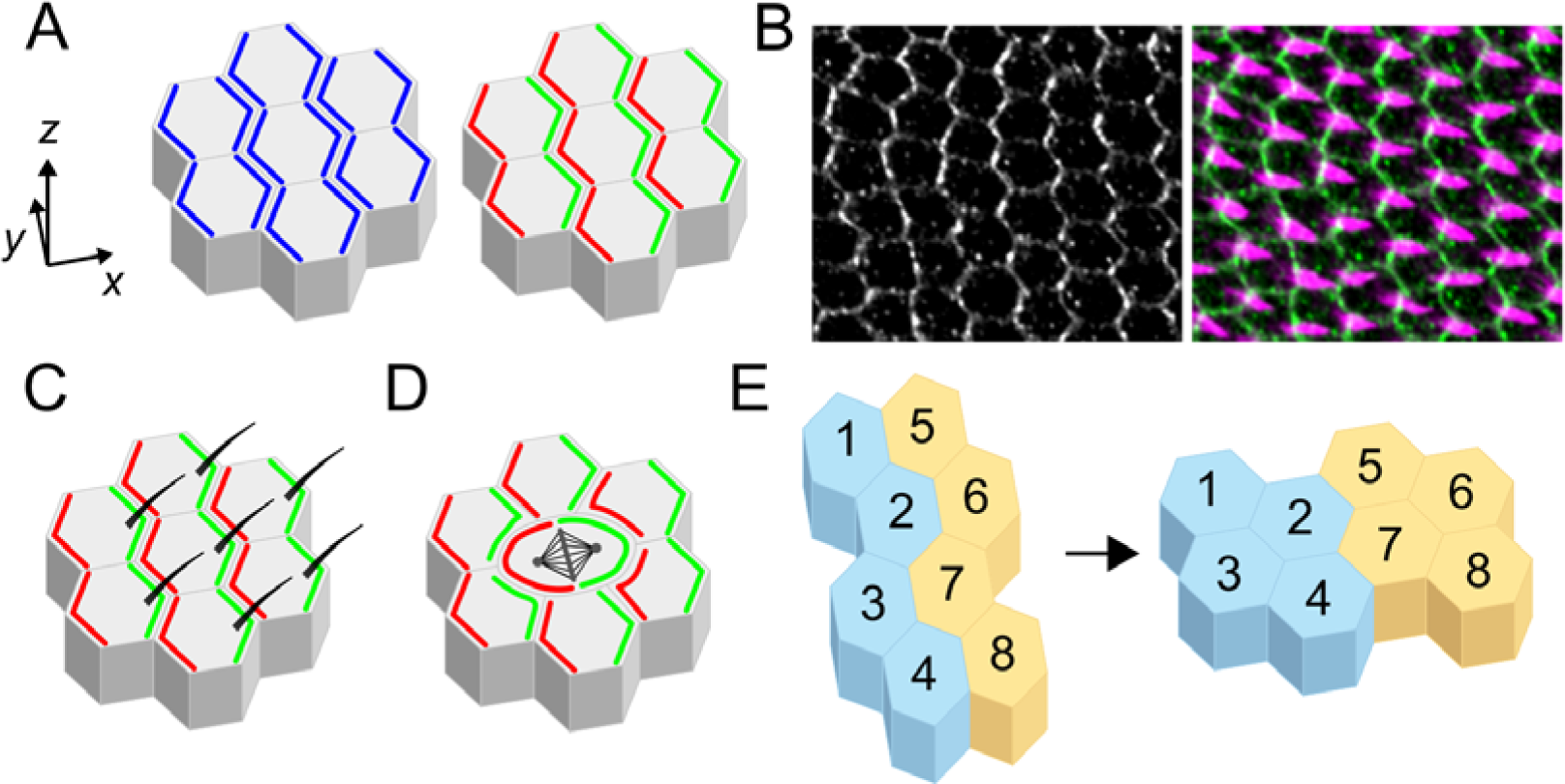
Planar polarity in epithelial morphogenesis. (**A**) In addition to polarising along an apicobasal axis (*z*), epithelial cells often exhibit planar polarity (also known as planar cell polarity) within the plane of the tissue (*x*, *y*). Planar polarity arises from the non-uniform distribution of polarity proteins, which may exhibit axial (enriched on opposite sides of each cell; blue) or vectorial (enriched on one side; red and green) polarity. (**B**) Wild-type *Drosophila* pupal wing (28h after puparium formation) stained for Vang (grey and green), which has vectorial polarity, and trichomes (magenta) (**C**, **D**, **E**) Planar polarity coordinates the alignment and organisation of cellular and multicellular structures. These include: the formation of hairs and bristles, such as the trichomes produced on the distal side of each cell on the adult *Drosophila* wing surface (**C**); oriented divisions, as observed for example in cells in *Drosophila* imaginal discs (**D**); and (**E**) polarised cell movements and rearrangements, such as during convergent extension.

This coordinated polarity can be readily visualised by the formation of oriented external structures such as hairs or bristles (Fig. 1B, C). It is also vital for fundamental functional roles that require cell coordination, such as oriented division (Fig. 1D) and convergent extension (Fig. 1E), thus disruption of these mechanisms results in disease [10]. Research into planar polarity establishment focuses on how long-range morphogen and mechanical gradients are interpreted at the cellular level [11], how cells communicate to coordinate information from upstream cues [12], and how downstream effectors alter cell behaviour and the forces underlying tissue formation [13].

Given the complexity of these processes, computational modelling plays an increasingly useful role in aiding our mechanistic understanding [14]. A key challenge is to interface models that include descriptions of cell shape, mechanics, and signalling on different scales. In this review, we consider the contribution of computational modelling first to planar polarity establishment, then to downstream mechanics, and the novel computational methods that study the interplay between them. For brevity, we consider animal tissues only, focussing primarily on *Drosophila* since the majority of planar polarity components have been extensively studied in that system.

## Modelling planar polarity establishment

Planar polarity can refer to any polarised protein or structure that breaks cellular symmetry in the plane of the tissue, occurring via multiple independent pathways. We begin by briefly summarising computational modelling of two key pathways: the Frizzled (Fz)-dependent or ‘core’ pathway, and the Fat (Ft)-Dachsous (Ds) pathway. We then describe the conserved anterioposterior (AP) patterning system active in the *Drosophila* embryonic epidermis.

### Core pathway

Components of the core pathway form asymmetrically localised molecular bridges between cells. The transmembrane protein Flamingo (Fmi; Celsr in vertebrates) can homodimerise via its extracellular domain across *inter*cellular junctions. Fmi interacts *intra*cellulary with two other transmembrane proteins, Fz and Van Gogh (Vang), which recruit several cytoplasmic factors (Fig. 2A). Since Fmi can homodimerise, it exhibits axial asymmetry (enriched on both sides of cells), whereas all other factors exhibit vectorial asymmetry (enriched on one side) (Fig. 1A). Fz and Vang appear to be the key components for recruiting other factors to apical junctional domains [15] and mediating cell communication of polarity [16,17], whereas the cytoplasmic proteins are thought to be responsible for polarity establishment [18-20] by amplifying initial asymmetries in Fmi, Fz and Vang through feedback interactions. The outcome of this pathway dictates, for example, the orientation of hairs on the *Drosophila* wing surface (Fig. 1B, C).

**Figure 2.**
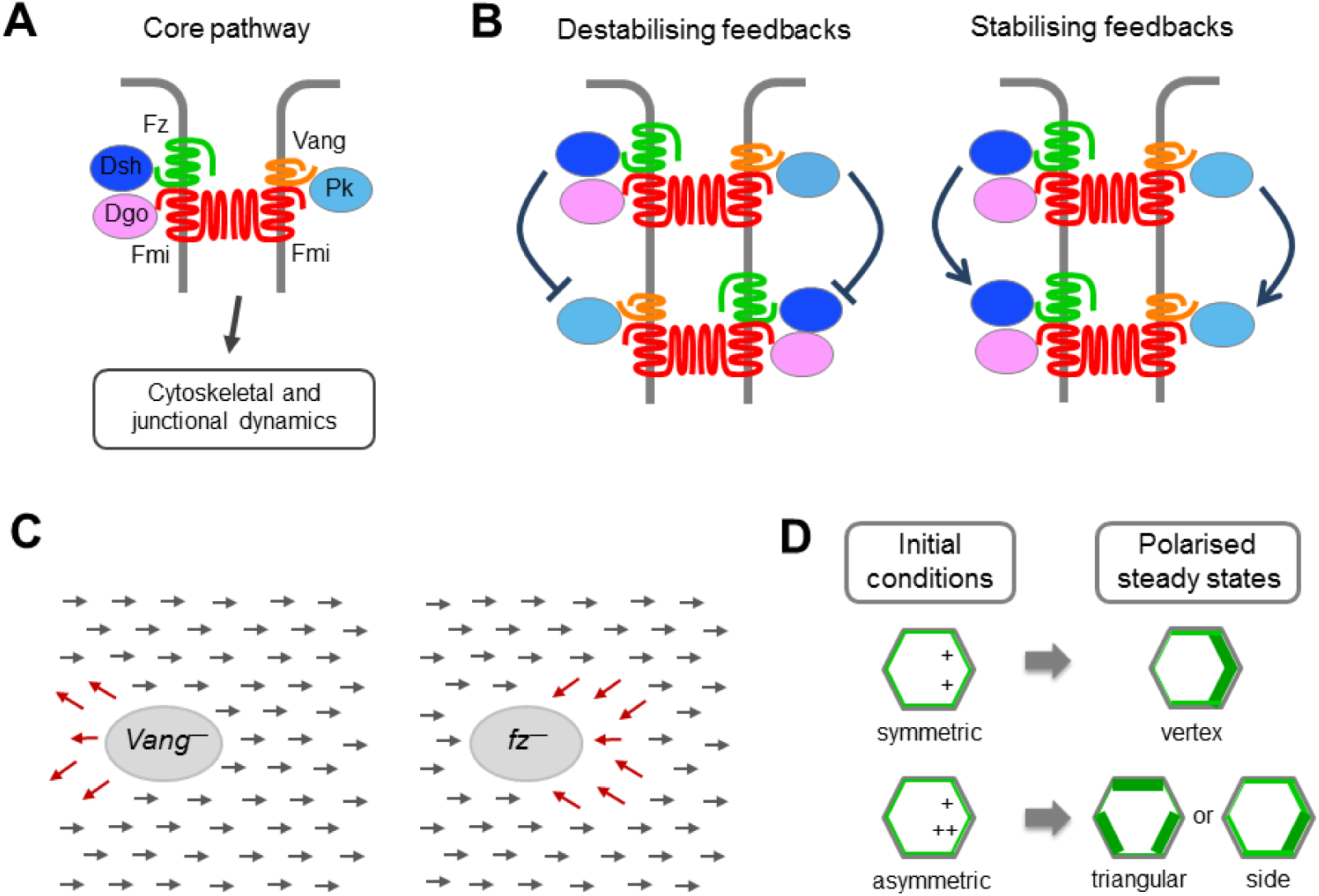
Computational modelling of the core pathway in *Drosophila* wing development. (**A**) Intercellular core protein complex arrangement at the adherens junction zone of *Drosophila* epithelial cells. The formation of an asymmetric intercellular complex involves the transmembrane proteins Frizzled (Fz; green) and Flamingo (Fmi; red) and the cytosolic proteins Dishevelled (Dsh; dark blue) and Diego (Dgo; pink) at the distal end of one cell, and the transmembrane proteins Vang Gogh (Vang; orange) and Fmi and the cytosolic protein Prickle (Pk; pale blue) at the proximal end of the adjacent cell. Polarised localisation of complex components leads to altered cytoskeletal and junctional dynamics, and thus altered cell mechanics. (**B**) Possible feedback interactions between non-transmembrane factors that, either alone or in combinations, could underlie amplification of asymmetry. For example, Dsh may inhibit Pk binding to Vang. (**C**) Schematic of non-autonomous phenotypes, observed in the *Drosophila* wing, around clones of cells mutant for Fz or Vang. (**D**) Schematic of 2D simulation results from Fischer et al [23], showing that the model of Amonlirdviman et al [22] does not give stable vertex polarised steady states in the absence of a persistent global bias. A uniform array of hexagonal cells is considered. In the upper panel, initial conditions are such that Fz is localised in all compartments of each hexagonal cell with a small initial bias (+) in the two distal compartments. This initial bias is amplified by the feedbacks, while symmetry is maintained, resulting in a final vertex polarity (thicker green edges). In the lower panel, an initial bias is applied but with a small difference (either+ or++) between the two distal compartments. Again the initial bias is amplified, but given the noise in initial conditions, vertex polarity is not maintained.

A variety of mathematical models have been proposed for the molecular wiring underlying this amplification [21]. In these models, asymmetric complexes form at cell junctions and feedback interactions occur between complexed proteins, such that either ‘like’ complexes of the same orientation are stabilised, or ‘unlike’ complexes of opposite orientation are destabilised, generating bistability (Fig. 2B). These models vary in complexity and include those based on Turing pattern formation mechanisms, using deterministic [22,23] or stochastic [24] reaction-diffusion approaches, and others based on the Ising model of ferromagnetism, which treat each cell as a ‘dipole’ that locally coordinates its angle with its neighbours [25]. Such models also vary in biological detail; from abstracted systems where two species bind to form a complex at junctions [26,27] to those including more defined molecular species. The latter necessitates many more kinetic parameters: for example, the model by Amonlirdviman et al [22] contains nearly 40 rate constants, diffusion coefficients and conserved concentrations whose values had to be estimated.

Domineering non-autonomous phenotypes, where a clone of cells mutant for a polarity protein influences the polarity of wild-type neighbours (Fig. 2C), have formed the basis for validating core pathway models at the tissue scale. Whether considering a one-dimensional row of two-sided cells [27], or a two-dimensional field of hexagonal [22] or irregularly shaped cells [28], various models are able to recapitulate these phenotypes. Importantly, modelling has bolstered our intuition on how polarity may be established and highlighted critical conceptual factors necessary for the system to work. For example, both the Amonlirdviman [22] and Le Garrec [24] models can generate tissue-level planar polarity when provided with a transient, rather than sustained, polarity cue; however, transient cues are not sufficient to ensure robustness of the resulting cellular polarisation (Fig. 2D) [23]. A number of biological candidates for a persistent global bias have been suggested, including the directional trafficking of Fz complexes along microtubules [29,30].

### Ft-Ds pathway

In contrast to the core pathway, there is strong evidence for a primary role of morphogen gradients in orienting the Ft-Ds pathway. In developing tissues, upstream morphogens specify opposing tissue gradients of Four-jointed (Fj), a Golgi-tethered kinase and Ds, a cadherin [31]. Ft and Ds are single-pass transmembrane proteins that can heterodimerise across intercellular cell junctions (Fig. 3A). They are both phosphorylated by Fj, which alters their ability to bind to one another [32,33]. Interestingly, although similar domains are modified on each protein, phosphorylation of Ft appears to improve its ability to bind to Ds, while phosphorylation of Ds is inhibitory. Work in *Drosophila* shows that Ft and Ds become asymmetrically localised within cells and that in turn recruits the atypical myosin Dachs to the distal side of cells [33-35]. Polarisation of this pathway can regulate tissue growth via the Hippo signalling pathway [36] and tissue shape by modulating tension at cell-cell junctions and orienting cell divisions [34,37,38], as well as coupling to the core pathway via the Pk isoform, Spiny-legs (Sple) [39].

**Figure 3.**
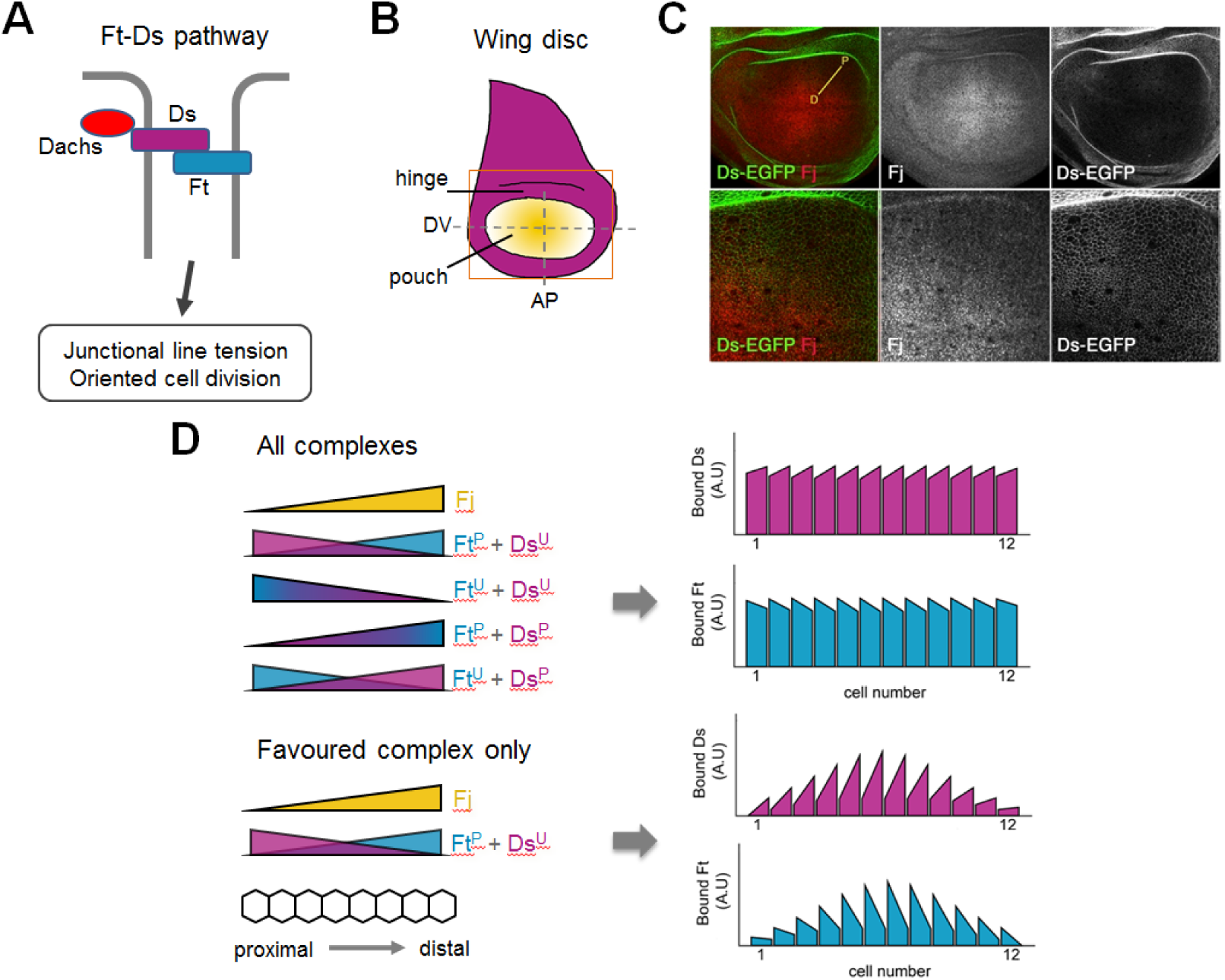
Computational modelling of Ft-Ds pathway establishment in the *Drosophila* wing. (**A**) Fat (Ft; turquoise) and Dachsous (Ds; purple) bind heterophilically to form asymmetric intercellular complexes. Dachs (red), an atypical myosin, is recruited to colocalise with Ds, where it modulates junctional tension and orients cell division. (**B**) Cartoon of *Drosophila* 3^rd^ instar larval wing disc. Ds (purple) is expressed at high levels in the hinge region, whereas Four-jointed (Fj; yellow) is expressed in a graded pattern in the pouch, which will go on to form the blade of the adult wing. Dorsoventral (DV) and anterioposterior (AP) compartment boundaries are shown by dashed lines. Orange box represents the cropped region shown in the upper panels of C. (**C**) Anti-Fj staining (red) of a wing disc expressing Ds-EGFP (green) as shown in Hale et al [42]. Fj is clearly graded along the proximodistal (PD) axis. (**D**) Simulation results based on the computational model of Hale et al [42]. Graded Fj leads to opposing gradients of phosphorylated Ft/Ds (Ft^P^, Ds^P^) and unphosphorylated Ft/Ds (Ft^U^, Ds^U^). Upper panel - all four possible heterophilic complexes form, listed in order of preferential binding (i.e. the top complex is the most favoured), leading to cellular asymmetry of bound Ft and Ds complexes that are largely uniform across the tissue. Lower panel - only the most favoured complex forms (Ft^P^ binding Ds^U^), thus polarisation and bound protein levels are much stronger in the middle of the tissue compared to the proximal and distal edges. Graphs show simulation results where each bar represents a cell, showing the relative amount of bound protein on the left and right sides in arbitrary units (A.U.).

While abstracted planar polarity models [8,26,27] could in principle be applied to the Ft-Ds system, models tailored to specific molecular interactions are limited. A recent phenomenological model examined the collective polarisation of the predominant complex – phosphorylated Ft (Ft^P^) binding unphosphorylated Ds (Ds^U^) – between cells in the *Drosophila* wing [40]. Either stabilising or destabilising feedback was found to amplify shallow graded inputs, but a combination of both more readily recapitulated experimental observations. By linking the strength of polarisation to a downstream tissue growth parameter, predictions were made and tested about the relationship between protein levels and overall tissue size.

Elsewhere, further molecular detail was included in a system of coupled ordinary differential equations describing interactions, again forming the predominant complex (Ft^P^ binding Ds^U^), in a one-dimensional row of cells [41]. However, for the majority of this study, the authors did not consider the orientation of those complexes at individual junctions, but only the asymmetry of total complexes across each cell, thus questions related to Ft and Ds polarity were not addressed. A more recent study used the *Drosophila* larval wing disc (Fig. 3B) to quantify the Fj gradient and Ds levels to initialise a one-dimensional reaction-diffusion model (Fig. 3C) [42]. Including all possible complexes of phosphorylated and unphosphorylated forms of Ft and Ds led to more uniform cellular polarity across the tissue (Fig. 3D). However, only considering the most favoured complex, as in previous models, resulted in greater variation in polarity and binding levels across the tissue. Coupled with experimental evidence, this supports the hypothesis that Fj acts on both Ft and Ds *in vivo*, but with opposing consequences, and illustrates the power of combining experimental and theoretical approaches in the same work.

### AP patterning system

In the *Drosophila* embryo, elongation of the body axis, known as germ-band extension, is driven by polarised cell movements and appears to occur independently of the core and Ft-Ds pathways [43]. Instead, evidence suggests that it is guided by striped pair-rule gene expression [44,45], although some contribution is also afforded to oriented cell divisions [46] and large-scale mechanical deformations [47]. The complex upstream gene-regulatory network consists of maternally derived morphogen gradients patterning gap gene expression, leading to stripes of pair-rule gene expression [48]. While the gap gene network has been extensively studied theoretically, uncovering shifting expression boundaries and the importance of transient dynamics of gene regulation [49,50], modelling of striped pair-rule gene expression and downstream processes remains limited.

## Modelling planar polarity pathway regulation of cell mechanics

The importance of mechanics in epithelial morphogenesis is well established [51]. Furthermore, increasing evidence suggests that a common role of planar polarity pathways is the spatial patterning of cell mechanics to affect consequent tissue-level morphogenetic processes such as convergent extension. Studies in both *Drosophila* and vertebrates reveal that downstream effectors include regulators of myosin II, actin and cadherins [52,53], which in turn affect anisotropy of local forces within an epithelial tissue (Fig. 4A). For example, the core planar polarity pathway has been implicated in polarised modulation of cell adhesion through trafficking of the adherens junction molecule E-cadherin. This appears to influence cell packing in the *Drosophila* wing and cell intercalation in the trachea [54,55].

**Figure 4.**
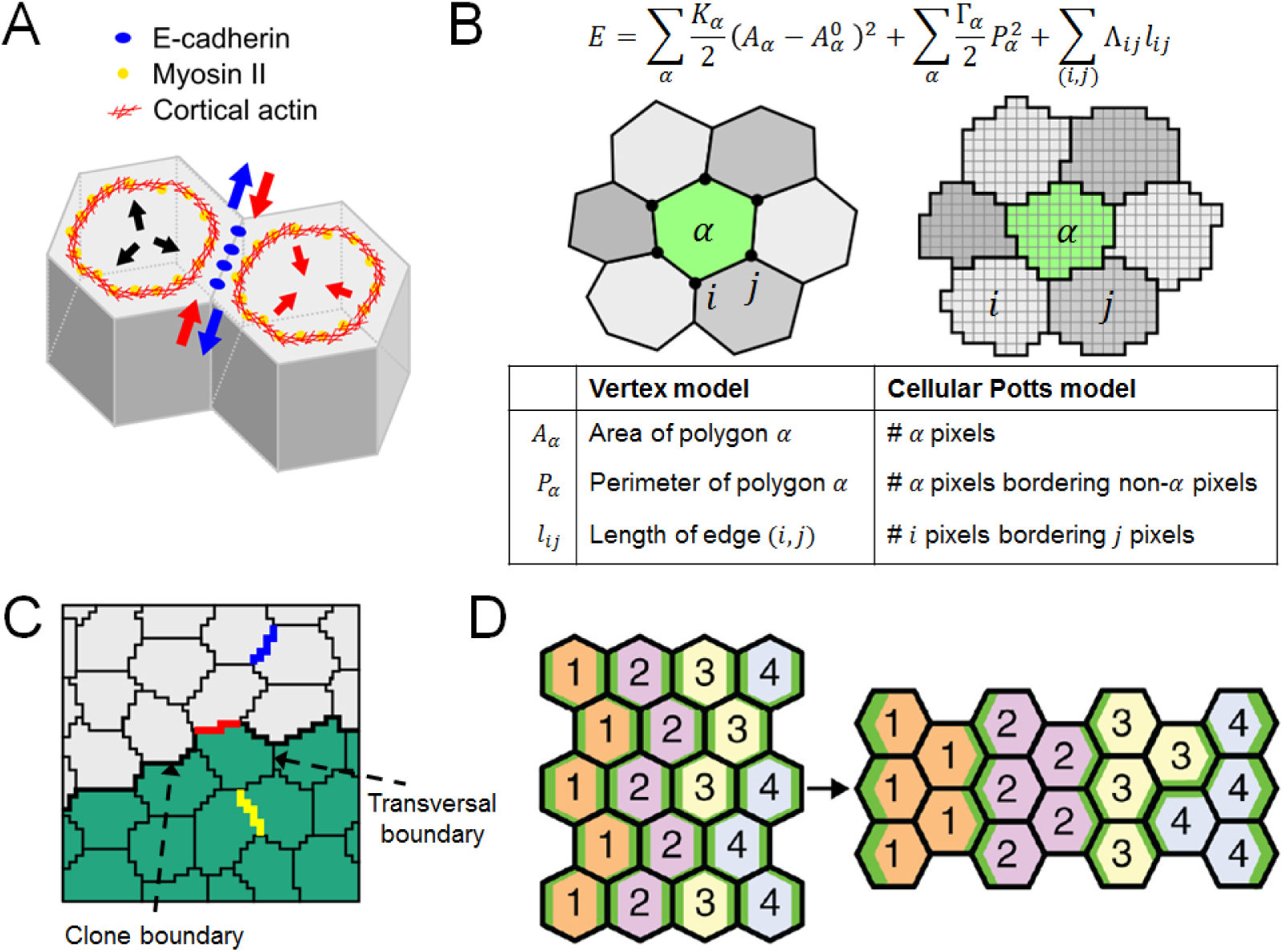
Computational modelling of the mechanics of planar polarised epithelia. (**A**) Schematic of the forces arising from apically localised adhesion molecules and cytoskeletal components in neighbouring epithelial cells. E-cadherin binding between tends to reduce surface tension and expand cell-cell junctions (blue arrows), while actomyosin imposes contractile forces at junctions and cell cortices (red arrows), the latter counteracted by intracellular osmotic pressure (black arrows). Each of these effector proteins can be regulated by upstream planar polarity signals. (**B**) Comparison of the vertex and cellular Potts models of epithelial dynamics. Either a force balance equation for each vertex (left) or Monte Carlo simulation and exchange of pixels (right) is used to drive the tissue toward a configuration of minimum ‘energy’,. (**C**) Cellular Potts model of somatic clone rounding in *Drosophila* pupal dorsal thorax [34]. Observed cell behaviours in Ft or Ds mutant clones are recapitulated by assuming that Dachs polarisation results in line tensions (Λ*_ij_*) taking a high value for cell-cell junctions at a clone boundary (red), an intermediate value for cell-cell junctions outside the clone (blue), and a low value for cell-cell junctions within the clone (yellow). (**D**) Vertex model of active cell intercalation during *Drosophila* germ-band extension [63]. Cell rearrangement results in stripes of cells of the same identity becoming adjacent. Myosin II is enriched at interfaces shared between cells of different identity (green). Convergent extension can be recapitulated by assuming that line tensions at cell-cell junctions (Λ*_ij_*) are increased by Myosin II enrichment and depend nonlinearly on the total length of contiguous interfaces a given cell has with cells of different identities, the latter assumption approximating the presence of actomyosin cables.

Nevertheless, models of polarity establishment typically assume that the dynamics of protein localisation occurs on a much faster timescale than cell shape changes, and thus consider a static cell packing geometry. To study dynamic cell shape changes requires coupling of models of planar polarity with tissue mechanics. To this end a variety of ‘cell-based’ models have been developed, which allow for the incorporation of cell signalling and feedback [56]. These include vertex [57] and cellular Potts [58] models, which approximate each cell's apical surface by a polygon whose vertices move according to a force balance equation, or a set of pixels that change stochastically to minimize an energy function, respectively (Fig. 4B). Each approach has its strengths and limitations [59]. Here we discuss a number of example studies.

### Core pathway

Inspired by evidence that the core planar polarity pathway can modulate cell mechanics, Salbreux et al [60] applied a vertex model to the ordered packing of cells in the zebrafish retina. Using a phenomenological differential equation model of planar polarity protein dynamics, the authors assumed that protein localisation modulates the ‘surface tension’ associated with cell-cell junctions and – through force balance – cell and tissue geometry. Geometry then feeds back on the localisation of planar polarity proteins. By comparing simulations under different hypotheses, the authors deduced that an extrinsic force (intraocular pressure) and progressive cell growth and division were required for the observed packing behaviour. Importantly, the authors tested model predictions by experiments with mutant fish such as those exhibiting increased intraocular pressure. Such work exemplifies the power of an approach in which experiments and computational models are tightly integrated.

### Ft-Ds pathway

As discussed above, the *Drosophila* Ft-Ds pathway is required for the planar polarisation of the atypical myosin Dachs. This in turn is required for orienting cell divisions during morphogenesis [37]. More recently a direct correlation between Dachs polarisation, membrane tension and tissue shape during growth has been made using a combination of modelling and mutant clone experiments in the *Drosophila* pupal dorsal thorax [38]. Following from earlier work linking Ft-Ds to mechanical control of morphogenesis [34], the authors explored why Ft or Ds mutant clones are rounded in shape, appearing to minimise their contacts with neighbouring cells, a process which is dependent on Dachs [37]. Notably, Dachs is enriched at clone boundary junctions and reduced at transversal junctions, those perpendicular to the clone boundary within the clone (Fig. 4C). This polarisation of Dachs correlated with altered line tension of these junctions. A cellular Potts model, with differences in tension at particular interfaces, was able to accurately recapitulate the clone circularity observed *in vivo*.

### AP patterning pathway

In the *Drosophila* embryo, the aforementioned pair-rule gene expression stripes lead to enrichment of Myosin II at AP borders and the adapter protein Bazooka/Par3 at dorsoventral (DV) borders [45,61], the latter recruiting E-cadherin to form adherens junctions. Planar polarisation of Myosin II, which drives the selective shortening of cell-cell junctions during active cell intercalation in germ-band extension [61], was recently discovered to be mediated by overlapping expression domains of Toll-like receptors [62]. This provides a combinatorial code where every cell along the AP axis has a different ‘identity’. To investigate how order is maintained as cells intercalate, Tetley et al [63] combined tissue-scale *in vivo* imaging and analysis with a vertex model incorporating differential junctional line tension between cells of different identities. Boundaries defined by polarised Myosin II, including parasegmental boundaries [47], were found to drive axis extension while at the same time limiting cell mixing. This work highlights the burgeoning recognition of the importance of ‘cables’ and other planar enrichments of actomyosin in coordinating morphogenetic processes. Future modelling efforts should include more mechanically explicit descriptions of how levels and polarisation of Myosin II and other effector proteins modulate cell mechanical properties. A pioneering example of such integration was recently proposed by Lan et al, who coupled modelling of polarisation of Rho-kinase, myosin and Bazooka with a vertex model, but restrict their attention to a relatively small number of cells [64].

## Interplay between mechanics and planar polarity

The above work seeks to understand the geometric and mechanical consequences of planar polarity signalling at the tissue level. However, recent evidence points to there being feedback, with adhesion and tension affecting tissue patterning pathways [13]. An extensive study used time-lapse imaging of *Drosophila* pupal wing development over several hours coupled with a vertex model showing that external tension elongates cells along the proximodistal axis and dictates the orientation of planar polarity [65]. Similarly, in the developing *Xenopus* embryo, mechanical strain has been shown to orient the global polarity axis [66]. Furthermore, in the mouse skin, Celsr1 symmetry appears to be broken by mechanical deformation along one axis [67]. Together, these results suggest a general mechanism where planar polarity proteins perdure on persistent junctions and are slow to accumulate on newly formed junctions allowing oriented cell rearrangements and tissue deformations to induce a new axis of asymmetry [11]. This further suggests that in some contexts core planar polarity polarisation is a passive process, governed by tissue-level changes. Conversely, the *Drosophila* Ft-Ds pathway is able to resist tissue strain and maintain its polarity in response to the graded signal of Fj, suggesting it is actively remodelled [39]. This is an intriguing area for future study where computational modelling may help to unravel why these pathways behave differently.

## Concluding remarks

We conclude by highlighting some extensions required to increase the utility of computational models in understanding planar polarity and tissue mechanics during development.

Several sources of biological complexity have not yet been incorporated or investigated within these models. A key consideration is the timescale over which a tissue can establish or remodel the asymmetric distribution of planar polarity components within a cell, versus the timescale over which mechanical changes occur. Notably, the rate of planar polarisation is likely to be strongly influenced by mechanisms such as directed vesicular transport and recycling of planar polarity components, but these have so far been neglected in current models. Furthermore, the significance of stochasticity and variability in polarity protein interactions and signal interpretation remain to be addressed, even though *in vivo* these are likely to contribute a significant degree of noise.

While two-dimensional computational models of patterned epithelial have established themselves as important tools, three-dimensional models remain limited and are typically restricted to imposed, static anisotropies in mechanical properties [68]. The extension of such models to allow for the dynamic simulation of planar polarity signalling remains to be tackled. For example, an intriguing link between core pathway planar polarity and three-dimensional tissue deformations was found by Ossipova et al [69], who demonstrated that planar polarity-dependent polarisation of the recycling endosome marker Rab11 is required for apical constriction and subsequent epithelial folding in the *Xenopus* neural plate.

Several software tools have recently been released for automated cell segmentation, tracking, and shape and polarity quantification in epithelial tissues [70-72]. This has coincided with the development of techniques to measure, infer, and manipulate forces *in vivo* [73,74]. Ongoing technical challenges associated with integrating the resulting data within computational models include developing efficient methods of simulating, and performing parameter inference and uncertainty quantification, on such models. Addressing these challenges will help to place computational models of planar polarity and tissue mechanics on a more quantitative footing, advancing their biological realism and power to guide future experiments.

## Acknowledgements

This work was supported by The Wellcome Trust (grant number 100986) and The University of Sheffield.

